# Inhibition of HIV-1 immune modulation by small molecules targeting viral Nef-host CD80 interface

**DOI:** 10.1101/2021.09.07.459239

**Authors:** Anusmrithi U. Sharma, Shweta Sharma, Gandhimathi Arumugam, Archana Padmanabhan Nair, Srinivas Ambala, Gurunadham Munagala, Kushalava Reddy Yempalla, Akankshi Munjal, Shreenidhi Rajkumar, Neelagandan Kamariah, Ashok R. Venkitaraman, Ramanathan Sowdhamini, Taslimarif Saiyed, Parvinder Pal Singh, Ram A. Vishwakarma, Satyajit Mayor, Anandi S. Karumbati

**Affiliations:** Centre for Chemical biology and Therapeutics, Institute of Stem Cell Science and Regenerative Medicine, GKVK, Bellary Road, Bangalore, India; National Centre for Biological Sciences, GKVK Campus, Bellary Road, Bangalore, India; Centre for Cellular and Molecular Platforms, GKVK Campus, Bellary Road, Bangalore, India; CSIR-Indian Institute of Integrative Medicine, Canal Road, Jammu-180001, India; Academy of Scientific and Innovative Research (AcSIR), Ghaziabad-201 002, India; The Cancer Science Institute of Singapore, Centre for Translational Medicine, National University of Singapore, Singapore 117599 & Agency for Science, Technology and Research (A*STAR), 8A Biomedical Grove, Singapore 138648

**Keywords:** immune modulation, HIV-1, Co stimulatory receptors-CD80/86, T-cell activation

## Abstract

HIV-1 causes diverse immunomodulatory responses in the host, including the down-regulation of co-stimulatory proteins CD80/86, mediated by HIV-1 protein Nef, blunting T-cell activation. Using a screening cascade of biochemical and cell-based assays, we identified potent small molecules representing three chemical scaffolds namely amino pyrimidine, phenoxy acetamide and bi-aryl heteroaryl carbamate which target the protein-protein interaction interface of CD80/86 and Nef with sub-micromolar potency. These molecules restore CD80/86 surface levels in HIV-1-Nef infected antigen presenting cells and T-cell activation. Nef-CD80 interface and small molecule binding sites were mapped by using computational docking and structural studies, followed by validation by mutational analysis. This analysis resulted in the identification of two key residues, K99 and R111, which were associated with down-modulation of CD80 surface levels by Nef and important for small molecule binding. Targeting these interacting residues disabled Nef-mediated down-modulation of CD80 surface levels, consequently restoring T-cell activation. Thus, we validate a new target, the Nef-CD80/86 protein-protein interaction interface, with a potential to develop new inhibitors to counteract the immunomodulatory consequences of HIV-1.

## Introduction

Over the past decade, there has been tremendous effort in finding newer therapies for HIV/AIDS. Increased use of anti-retroviral drugs has been accompanied by the steady increase in HIV drug resistance and viremia that often result in immunosuppression leading to morbidity^3^. Drug resistance is mainly transmitted at the time of infection or acquired during previous treatment for instance in women given anti-retroviral drugs to prevent mother-to-child transmission of HIV^4^. Other treatment strategies like using anti-HIV-1 antibodies (bnAbs) are reported in combination with ART which have the capacity to impact on HIV-1-specific T cell immune responses in infected humans but whether it controls the virus remains to be determined^5^. In addition, with the vaccine trials failing to elicit broad plasma neutralization of primary virus isolates^6^, there is an urgent need to fast-track the transition to newer antiviral drug regimens which needs to be administered in multi-drug combinations in order to combat the ever-evolving virus. Increasing drug resistance also emphasizes the requirement of new molecules that can target the host-viral interface with high specificity^7, 8^.

To obtain targets for the next generation of HIV-1 therapy we explored the interaction interface of the essential HIV-1 accessory protein, Nef^9^, with host proteins. Nef is a 27-35 kDa protein expressed during early phases of viral replication and helps maintain a constant state of infection by disrupting T-cell activation, thereby allowing evasion of the host immune system^10,11,12^. Evidence supporting a direct role for Nef in HIV disease comes from transgenic mouse models developed, in which a CD4-derived promoter was used to express Nef in various tissues including thymus, kidney and lungs which progressed into AIDS-like phenotype, featuring CD4+ T-cell loss, thymic involution, splenic atrophy and subsequent kidney and lung pathology^13^. This phenotype in many aspects’ mimics human AIDS. Also in patient populations, fortuitous deletions in the *nef* gene in HIV infected patients of the Sydney Blood Bank Cohort remain essentially free of AIDS related symptoms^14, 15^. In another study, the proportion o*f nef* gene defects was found to be significantly higher in Long-Term Non-Progressors (LTNPs) compared to progressors ^16,17,18^. These animal and patient studies suggests that Nef plays a pivotal role in pathogenesis and AIDS-like progression in HIV-infected individuals^19^.

Nef functions by re-engineering the levels of many surface proteins such as MHC-I, CD4, CD28, CXCR4, and CD3 in infected cells, and redirecting them to endosomes^20, 21^. Nef interacts either directly or indirectly with multiple host partners and functions to increase the pathogenesis of the virus^22, 23^. Our previous work has highlighted interactions of Nef with the host cell surface co-stimulatory proteins CD80 and CD86^24, 25^. Down-regulation of CD80/86 is sufficient to cause impaired naïve T-cell stimulation *in vitro* and *in vivo*^26^. We have shown that by administering just the cytoplasmic tails of CD80 and CD86 in Nef-expressing cells, CD80/86 down modulation is prevented, making this host-viral interface amenable for developing new chemical biological tools and providing targets for therapeutic intervention. The variety of functions carried out by Nef is based on its ability to interact with multiple cellular proteins. This is possibly due to its structure; it has many flexible regions, a feature unusual for cytoplasmic proteins. The unstructured regions of Nef may ease the allosteric adjustments required for interaction of Nef with different proteins. However, the flexible and unstructured features of Nef protein add substantial challenges to structurally characterize the full-length Nef protein and study their binding sites.

In the current study, we show that Nef directly binds to cytoplasmic tail peptides of CD80 and CD86. We have identified small molecules, which can abrogate Nef interactions with CD80 and/or CD86. The compounds mainly belong to 3 scaffolds amino pyrimidine (**AP**), phenoxy acetamide (**PA**) and bi-aryl heteroaryl carbamate (**BC**) having nanomolar to micromolar inhibition potencies *in vitro*. Representative actives from these scaffolds were then validated in functional cell-based assays for reverting the down regulation of cell surface Nef-mediated co-stimulatory protein expression and re-establishing T-cell activation. These identified actives also reversed similar Nef-mediated effects after viral infections.

To further improve the efficacy of these leads, we used an *in-silico* approach to explore the binding mode of Nef with the co-stimulatory molecules. Full length Nef protein was modeled and potential binding sites for CD80 were identified. Based on these predictions a selected subset of CD80 interacting residues in Nef were mutated and the corresponding mutants were examined in assay platforms to confirm the model and its functional consequences. The understanding of the druggable pocket opens the scope for future lead optimization work. Altogether, we report a chemical strategy to inhibit Nef-mediated immunomodulatory functions, which prevents immune evasion of HIV-infected cells. While these molecules may be developed for future therapeutic intervention, at the current stage, they may be used as chemical biology tools to understand the role of host-pathogen interface in the form of Nef-CD80/86 interaction surface in HIV immune evasion.

## Results

### Identification of potent Inhibitors that disrupt Nef-CD80/CD86 Interaction

In our previous work, we have shown that cytoplasmic tail peptides from CD80 and CD86 compete with the down modulation of surface CD80/86 by Nef protein in cell based assays^25^. For our assays we have used Nef F2 recombinant protein of Subtype-C origin (the alignment of the Nef sequence from this subtype with the commonly used subtype B is presented in **Sup. Fig. S1a**). To ascertain direct binding of Nef to the cytoplasmic tail peptide of the co-stimulatory receptors CD80 or CD86 in microscale thermophoresis (MST) assay^27^, we added 20-mer peptide of CD80 or CD86 (**Fig. 1a, Sup. Fig. S1b**) to fluorescently labelled Nef. A 16-point titration of peptides against Nef was carried out (see details in Star Methods). Nef-CD80 interaction showed a saturation curve with a kD of 27 µM (**Fig.1b**). Nef-CD86 interaction showed a sigmoidal curve with 112 nM kD, indicating a higher affinity interaction (**Fig.1c**). Having verified the direct binding of the CD80 and CD86 peptides with Nef in MST, we immobilized CD80/86 cytosolic peptides in a microtiter plate and examined the binding of Nef in an ELISA assay (Scheme, **Sup. Fig. S1c**). There was a 4-fold increase in the fluorescent signal due to binding of Nef to the 20-mer CD80/86 cytosolic tail peptides compared to wells with an unrelated peptide (from CD74), or with no peptide controls (**Fig. 1d**). We used this ELISA format to screen and identify inhibitors of Nef-CD80 and Nef-CD86 interaction.

**Fig. 1.**
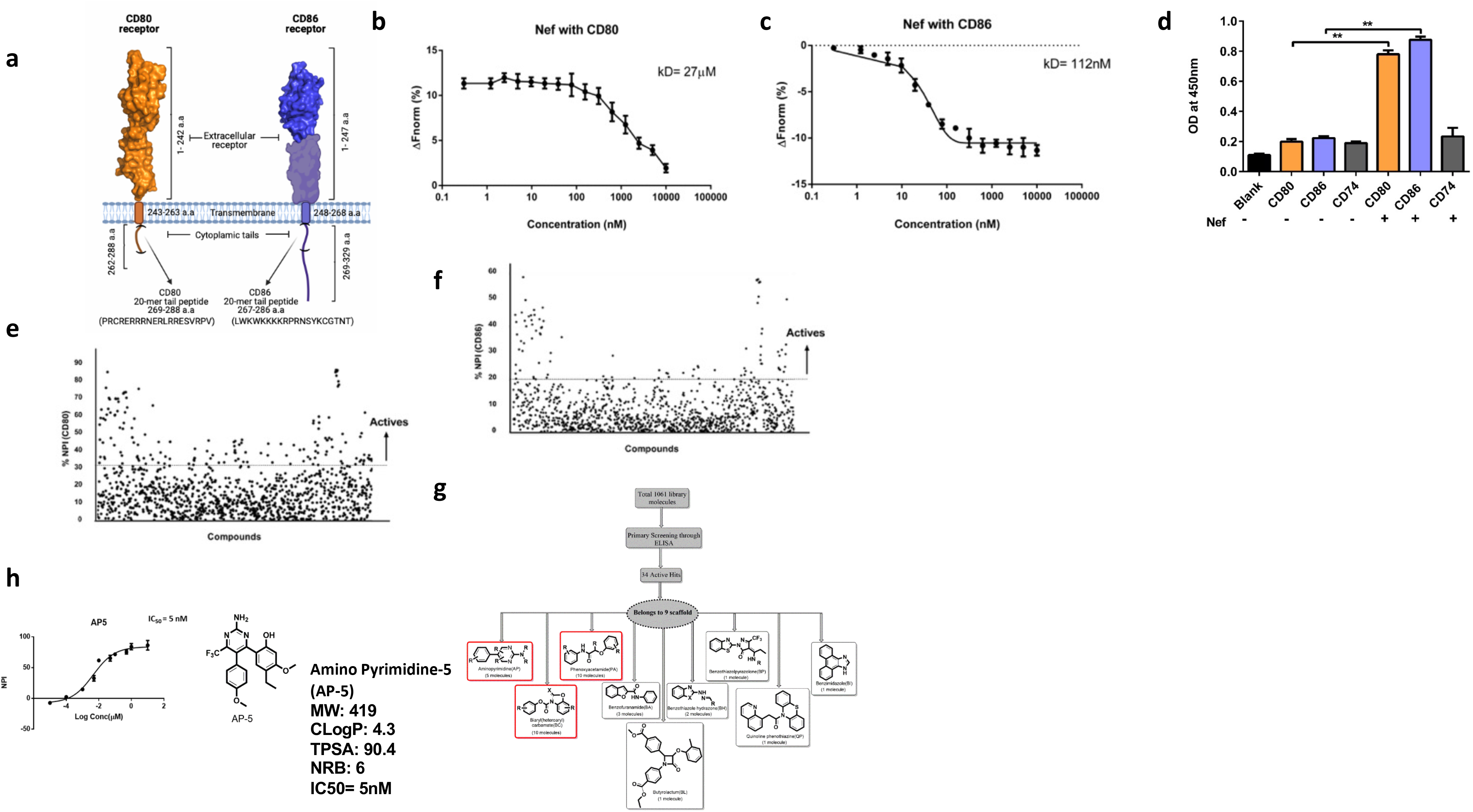
Nef directly interacts with the cytoplasmic tail peptides of CD80/86. **a)** Illustration shows the CD80 and CD86 receptors with their extracellular, transmembrane and cytosolic tail region marked. The 20-mer peptide region of CD80 (PRCRERRRNERLRRESVRPV, 269 to 288 a.a) and CD86 (LWKWKKKKRPRNSYKCGTNT, 267-286 a.a) is highlighted within the cytoplasmic tail domain (**b)** Graph shows direct binding of Nef to CD80 as measured by Microscale scale thermophoresis (MST). CD80 peptide was titrated from 950 µM in 2-fold dilutions upto 16 points against a fixed Nef concentration (35 nM) in the final reaction volume. A curve with upper saturation with kD=27 µM was obtained; x axis= peptide concentration (nM) and y axis= percentage normalized fluorescence (ΔFnorm); Plots represent the mean ± SD (error bars) from three independent experiments **(c)** Similarly, CD86 peptide was titrated from 10 µM with a 2-fold serial dilution upto 16 points against a fixed Nef concentration (35 nM). A sigmoidal curve with kD=112 nM, showing higher affinity as compared to CD80. **(d)** Graph shows ELISA assay where change in OD is observed when the immobilized CD80 and CD86 cytosolic peptides binds to Nef. The OD measurement was done at 450nm. CD74, a negative peptide control shows minimal OD value **(e)** Graph shows Normalized Percentage Index (NPI) on a normal distribution curve for statistical significance of active compounds across qualified plates showing CD80 actives. X axis= the number of compounds screened in ELISA assay; y axis= normalized percentage Inhibition of compounds; of Z-factor>0.5 analysis was used to qualify the plates. Compounds with NPI>30% for CD80 was considered as hits **(f)** Similarly, NPI normal distribution curve for CD86 with Cutoff percentage for CD86 NPI>20% was considered as hits **(g)** Scheme shows the hits belonging to 9 scaffolds that were identified in the primary screen **(h)** Dose response curve of a hit compound from “AP” scaffold; x axis= log concentration of compounds; y axis= Normalized percentage Inhibition, (inset) Structure of AP5 compound and its molecular properties

Small molecules were hand-picked for screening based on their drug-likeness and prior knowledge of potential protein-protein disruptors^7^. A threshold value was set to select actives with CD80≥30% and CD86≥20% cut-off for the Normalized Percent Inhibition. Based on this criterion, 33 actives belonging to nine scaffolds were identified from the screen (**Fig.1e, f**). **Fig.1g** depicts flow diagram of step-wise filtering process. After eliminating singleton scaffolds; we identified hits that belonged primarily to three scaffolds namely amino pyrimidine (**AP**), phenoxy acetamide (**PA**) and bi-aryl heteroaryl carbamate (**BC**), which comprised of 20 compounds (**Supp Table 1**), and these were selected for resynthesis (**Sup. Fig. S2**). Compounds from all three scaffolds showed half maximal Inhibitory Concentration (IC_50_) in nanomolar ranges (**Fig. 1h; Supp Table 1**). Two independent experimental dose response data correlated very well with an R^2^ value of 0.9 (**Sup. Fig. S3**).

In summary, the biochemical screen identified more interaction inhibitors of Nef-CD80 than Nef-CD86, which correlates well the MST binding isotherms indicating that the Nef-CD86 interaction is of higher affinity than the Nef-CD80 interaction. One representative compound from each scaffold with high potency against Nef-CD80 interaction was chosen for further cell-based assays.

### Small molecules block Nef-mediated internalization of cell surface CD80/CD86 receptors

Before starting efficacy studies, we evaluated the cytotoxicity of the compounds (**AP5**, **PA4** and **BC5**), as measured by WST-1 (Water Soluble Tetrazolium-1) assay. The cytotoxicity index at the highest concentration of 100 µM for 24 h was around 17% for **AP5** and **PA4** whereas **BC5** was around 20% (**Sup. Fig. S4**). The maximum concentration tested was 100 µM, and the concentrations chosen for the efficacy work were mostly non-toxic for the assay durations.

The cellular efficacy assay was developed based on observations from our previous work: the delivery of cytoplasmic CD80/86 tail peptides into the cell cytoplasm was able to compete with and abrogate Nef-mediated internalization of cell surface CD80/86^25^. We expected the small molecules identified by the biochemical screen to behave in similar fashion to the cytosolic peptides. Surface levels of CD80/86 receptors in cells in culture with or without Nef protein were determined, and we observed a significant loss of surface levels of CD80 and CD86 in the presence of Nef in at least 3 types of Antigen Presenting Cells (APCs), including monocytes (**Fig. 2a**), consistent with our previous work^25^. The loss of CD80/86 surface receptors was not observed with delivery of other non-specific proteins such as ovalbumin and β-lactoglobulin which confirms that the internalization of surface receptors CD80 and CD86 was indeed due to Nef (**Sup. Fig. S5**). We used RAJI, a B-lymphocyte cell line for our experiments since this cell line has high levels of CD80 and CD86, and chose to focus on targeting on the Nef-CD80 interface.

**Fig. 2:**
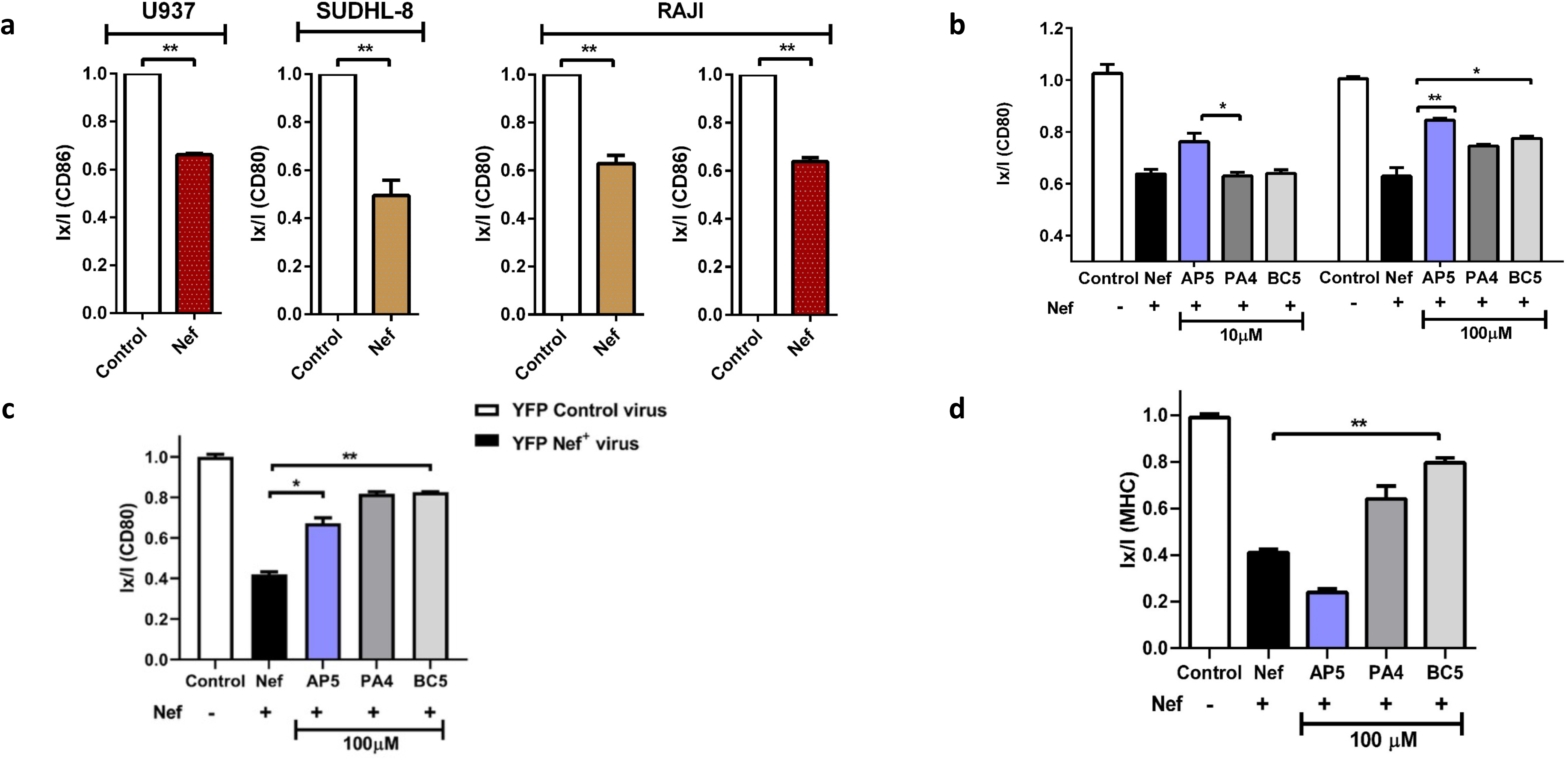
Cell based assay screening of active compounds from ELISA. **(a)** Nef mediated down-regulation of surface CD80 or CD86 in 3 different cell lines as indicated. FACS data showing the normalized surface levels of CD80; I_x_/I (y axis) where I_x_ is the average fluorescence intensity in the indicated condition (from a triplicate) and I is the Median of normalized negative control (No Nef control) **(b)** FACS data shows restoration of CD80 receptors in RAJI cell line after pre-treatment with 3 representative compounds **AP5, PA4 and BC5** at 10 and 100 µM for 24 h and analysis after 2 h post Nef protein delivery **(c)** RAJI cell line was infected with viral particles (Nef-YFP and YFP alone control cells) in viral infection assay and surface CD80 receptors with compounds were measured by flow cytometry **(d)** Effects of inhibitors on Nef-MHC-I interactions. RAJI cells were treated with compounds at 100 µM and then stained with anti-MHC-I antibody. MHC-I was detected by flow cytometry and shown as I_x_/I plots. Compound **AP5** shows no restoration of MHC-I indicating its specificity for the Nef-CD80 interface.

To test the effect of the compounds on Nef-mediated CD80 down modulation, cells were pre-treated with the **AP5**, **PA4** and **BC5** at two concentrations 10 and 100 µM for 1 hour. Purified Nef protein was delivered into the cells and surface CD80 levels were analyzed by flow cytometry. **AP5** inhibited Nef-mediated down modulation of CD80 at both 10 µM (*p ≤0.05) and 100 µM (**p ≤ 0.01) thereby restoring CD80 levels. Compounds **PA4** and **BC5** also showed a significant restoration (*p ≤0.05) of surface CD80 levels only at 100 µM (**Fig. 2b**).

We further validated the ability of these three compounds to reverse the effects of Nef in RAJI cells transduced with Nef-containing virus (YFP-tagged Nef) or Nef-deficient control virus (expressing only YFP). RAJI cells were pre-treated with **AP5, PA4** and **BC5** for 24 hours’ prior-exposure to virus and further incubated with virus for 96h and then surface levels of CD80 receptors were estimated by flow cytometry, there was ∼50% loss of CD80 surface receptors in cells transduced with Nef-containing virus as compared to control Nef-deficient virus infected cells. In cells pre-treated with compounds, all 3 compounds significantly reversed Nef-mediated internalization of CD80 receptors at 100 µM (**Fig. 2c**).

### Compound AP5 is a selective inhibitor for Nef-induced CD80 down modulation

Since Nef also downregulates other cell surface molecules such as MHC-I and MHC-II in APCs and CD4 in T-cells, apart from CD80/86 during viral infection, we determined the effect of these compounds on the levels of MHC-I^28^ in Nef-transduced APCs. After compound treatment, we observed that **PA4** and **BC5**

reversed Nef-mediated effect of MHC-I down modulation while **AP5** did not alter Nef-mediated down regulation of MHC-I even at 100 µM concentration (**Fig. 2d**). Together, these data suggested that while all compounds disrupted Nef interactions with CD80 and prevented the down modulation of CD86, the Nef-activated cascade that triggers MHC-I down modulation was reversed only by **PA4** and **BC5.** This indicates a very specific role for **AP5,** and a more broad-spectrum role for **PA4** and **BC5**.

### Restoration of T-cell activation in virus infected APCs by inhibitors

The co-stimulatory receptors CD80/86 along with MHC-I are needed for T-cell activation. Previously it was shown that HIV-1 Nef-CD80/86 interaction impairs naïve T-cell activation in *in-vivo* and *in-vitro* mouse systems^24^. We therefore ascertained if the small molecule inhibitors were able to reverse Nef-mediated T-cell inactivation. A co-culture assay system was adapted from a previously reported cell-based assay for testing CD80 inhibitors^29^ (**Fig. 3a**). Briefly, a non-replicative retroviral vector with Nef transgene (**Sup** **Fig. 6** **shows a** comparison of the non-replicative retroviral vector with infectious HIV-1) was transduced in APCs to reduce CD80/86 levels at the surface . These APCs were then co-cultured with T-cells in the presence of anti-CD3 antibody and functional T-Cell activation was assessed by measuring IL-2 levels; relevant controls included T-cells alone, T-cells and B-cells in the absence of anti-CD3 antibody (**Fig. 3b**).

**Fig. 3:**
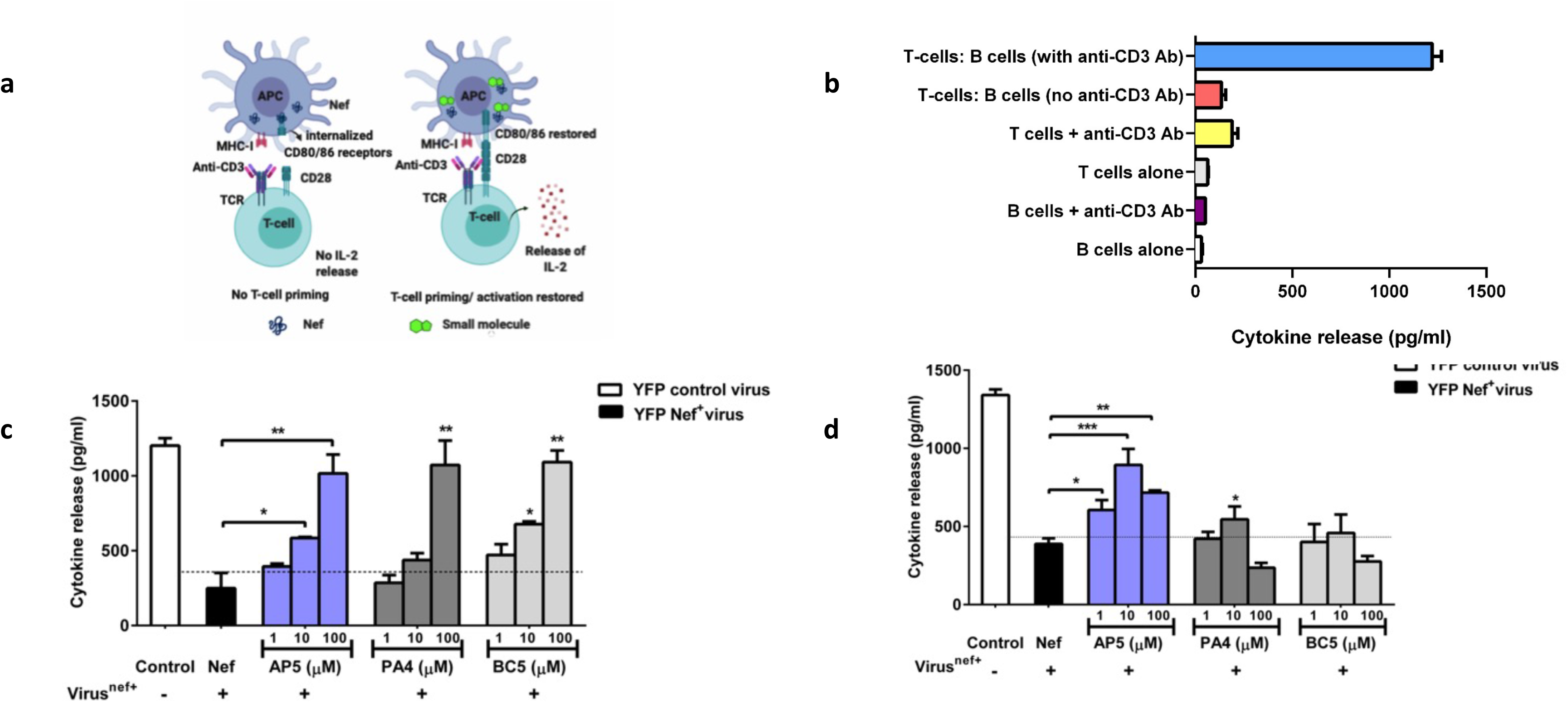
Restoration of functional T-cell activation in a viral infection assay. **(a)** Schematic of a functional assay for screening of compounds that disrupt Nef-CD80/86 interactions in a virally-infected cell. Functional T-Cell activation is based on APC-T cell co-cultures. The APC has CD80/86 and concurrent presence of anti-CD3 antibody promotes T-cell activation (as measured via IL-2) in co-cultured T-cells **(b)** Graph shows cytokine release in functional T-cell activation assay where antigen presenting cell RAJI (B-cells) and Jurkat-cells (T cells) were co-cultured as indicated (with/without anti-CD3 antibody); T-cells alone, B-cells alone controls do not show measurable IL-2 release. **(c)** Graph shows quantification of cytokine (IL-2) released after T-cells and B-cells co-culture for 3 h; here B-cells were pretreated with the indicated concentrations (1, 10, 100 µM) of compounds for 24 h followed by viral infection for 96 h **(d)** Graph shows quantification of cytokine (IL-2) released after T-cells and B-cells co-culture for 3 h; here B-cells were first infected with virus for 96h and then treated with compounds at 1, 10, 100 µM for 24 h. IL-2 release (pg/ml) was determined by ELISA by plotting against an IL-2 standard curve. Note: viral infection reduces IL-2 release, and all 3 compounds showed a dose dependent restoration of IL-2 release. **AP5** showed IL-2 release at 1 µM.

We tested two different modes of addition of compounds: in the first mode, we pre-treated APCs with 1, 10 and 100 µM of compounds (**AP5**, **PA4** and **BC5**) for 24h then exposed the cells to Nef-carrying viral particles and assayed for IL-2 release. We observed a dose dependent response with all 3 compounds **AP5, PA4** and **BC5** showing a 4-fold increase (**p ≤ 0.01) in IL-2 release at 100 µM as compared to Nef virus control. At lower concentrations, compounds **AP5** and **PA4** showed 2-fold increase (*p ≤0.05) at 10 µM. **PA4** did not show significant IL-2 release at 10 µM (**Fig. 3c**), thus, indicating restoration of T-cell activation in a dose dependent manner.

In the second mode, we added compounds at 1, 10 and 100 µM concentrations post viral exposure for 96h. **AP5** showed significant increase in IL-2 levels (3-fold) at all concentrations when compared to Nef virus control (**p ≤ 0.01), while **PA4** showed 2-fold increase (*p ≤0.05) in IL-2 levels at 10 µM concentration and no changes in IL-2 levels were observed in **BC5**. Interestingly, there was a reduction in IL-2 release at 100 µM dose in comparison to 1 µM and 10 µM, perhaps due to the toxicity associated with this dose alongside viral effects (**Fig. 3d**). Thus far, these data shows that **AP5** is a potent molecule both *in vitro* biochemical assays and in restoring T-cell activation in an assay designed to assess the role of co-stimulation dependent T-cell activation mediated by CD80/86.

### Structural insights into Nef-CD80 interaction

To further understand the nature of inhibition of Nef-CD80 interaction by small molecules such as AP5, we chose to analyze the structure of Nef-CD80-interaction pocket. This requires the characterization of the interaction surface between Nef-CD80 and an analysis of the ligand binding pocket for CD80. Despite repeated attempts we were unable to obtain crystals of full-length Nef that diffracted better than 4 Å, wherein the structure could be fully resolved. In the absence of a high-resolution X-ray structure of the full-length Nef, we created a computational model of Nef using a multi-template modeling approach. The major structural information was acquired from the NMR structure PDB ID: 2NEF, as this structure has information for the highly flexible loop region of the core domain (55-66) which contains important interacting residues^30^, as well as from the crystal structure PDB ID: 3RBB which contains structural information of C-terminal folded core (residues 79-206). After modeling, the lowest energy state structure was obtained by energy minimization via SYBYL (Version 7.1) (Tripos Associates Inc.) and validated using PROCHECK. PROCHECK results for the model shows more than 95% of the residues are in allowed regions (79.9% in the strictly allowed region and 17.2% in partially allowed region of the Ramachandran plot) which is better than the template structure (62.3% in the strictly allowed region and 34.2% in partially allowed region of the Ramachandran plot).

The structure of full-length Nef can be divided into two parts: a flexible and structurally diverse N-terminal region of about 70 residues followed by a well- conserved and folded core domain of about 120 amino acids. The core domain is the only part of the Nef protein which has a stable tertiary structure. It forms an α-β domain in which a central anti-parallel β-sheet of four strands (β1- β4) is flanked by two long anti-parallel α helices (α4 and α5) and two short α helices (α1 and α5). Residues 60-71 and 149-180 form flexible solvent exposed loops^19^ (**Fig 4a**).

**Fig. 4:**
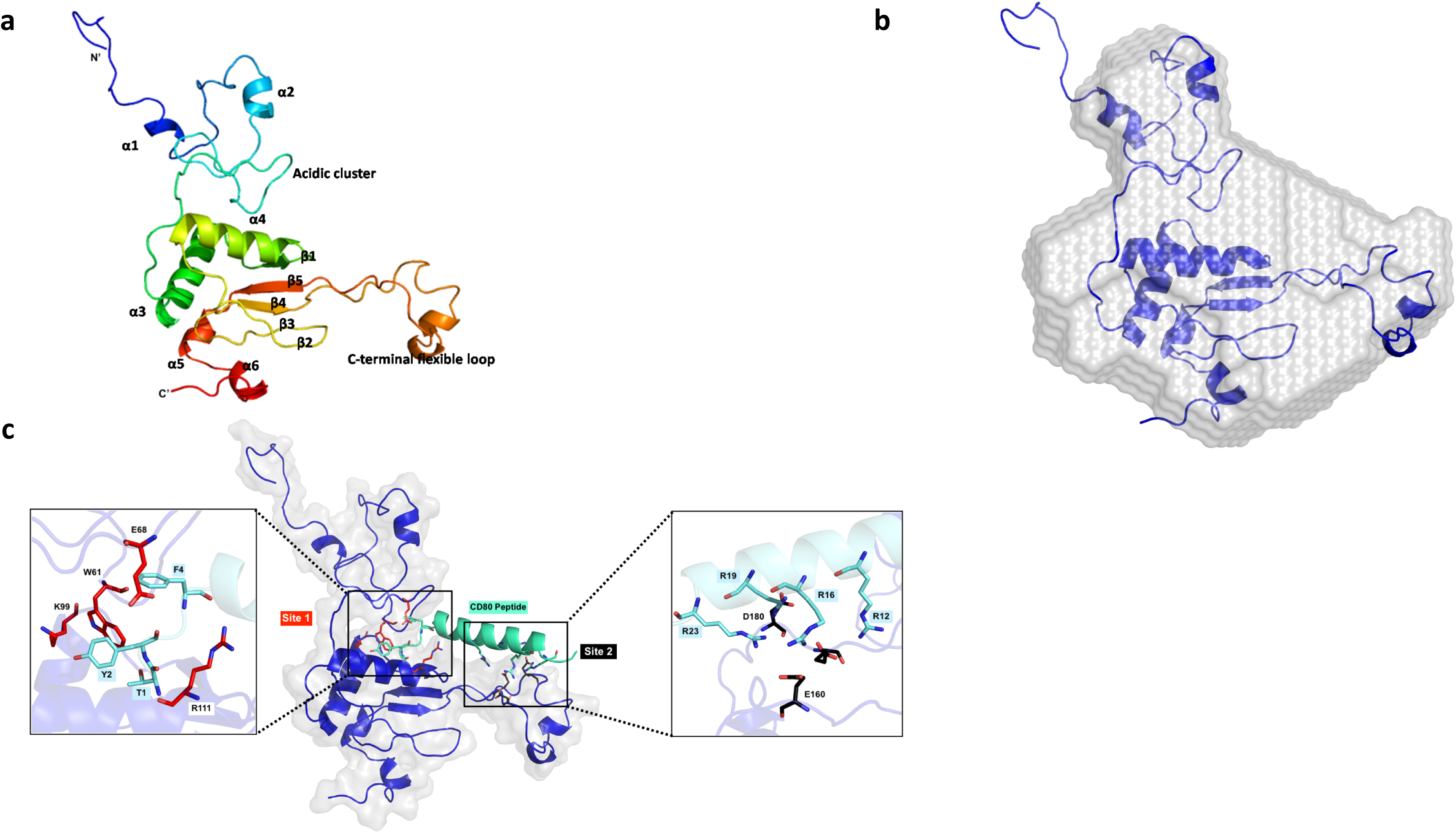
Modelling of Nef with multi-template computational approach. **(a)** Cartoon representation of the predicted structure of Nef shows flexible N-terminal region and well-conserved core domain, colored in accordance with their position (N-terminal in blue to C-terminal in red) with respective α-helices and β-sheets as indicated. **(b)** The *ab initio* shape of the solution structure of the Nef (blue color) from the SAXS data (grey surface) fits well with the computational model (cartoon representation) **(c)** Surface representation of HIV-1 Nef is depicted with the best CD80 binding pose. CD80 peptide in a docked pose (cyan color) in Nef obtained using SiteMap program. The inset shows the important residues of Nef involved in interaction with CD80 at Sites 1 and 2.

An independent verification of some aspects of the model was obtained from the predicted Small Angle X-ray Scattering (SAXS) envelope of soluble Nef protein. SAXS-patterns of full-length Nef were obtained at three different concentrations 1, 3 and 5 mg/ml (**Sup. Fig. S7a**). The Guinier plots at low angles appeared linear and confirmed good data quality with no indication of protein aggregation (**inset Sup. Fig. S7a**). The derived *R_g_* values and the calculated maximum particle dimension (*D_max_*) values were reported. The *R_g_* values extracted from the *P(r)* function are in agreement with the *R_g_* values extracted from the Guinier region (**Sup. Table 2**). The estimated *R_g_*, *D_max_* and molecular mass of the full-length Nef suggest a concentration dependent increase in the *R_g_*, *D_max_* and molecular mass values observed with Nef addition. Visual inspection of the normalized Kratky plot reveals significant deviation from a bell-shaped profile which depict an inherent structural flexibility of Nef (**Sup. Fig. S7b**). The averaged solution shape calculated using the 1 mg/ml scattering data clearly indicated that Nef is monomeric in solution. This solution model also revealed a two-domain architecture, a large domain that is well overlaid with the available 3D structure of folded C-terminal core (PDB ID: 3RBB) (**Sup. Fig. S7c**)^31, 32^. The small domain corresponds to the N-terminal region (residues 1-78) that contain a long flexible loop (residues 24-68). While structural details of N-terminal region are available from NMR studies of a peptide regions from residues 2-26^33^ and 2-57^30^, information about the relative orientation with C-terminal core domain is missing. The SAXS data envelope along with the computational model (**Fig. 4b**) provides structural information about spatial arrangement of the C-terminal folded core and the flexible N-terminal region of full-length Nef in solution, consistent with the computational predictions.

### Identification of crucial residues involved in Nef-CD80 interaction surface

We next utilized the computational prediction of full-length Nef structure and molecular docking studies with the cytoplasmic tail of CD80, to identify key residues at the interaction surface. The putative binding sites of cytoplasmic CD80 to full length Nef were mapped onto the template model utilizing SiteMap program for binding site prediction. In characterizing binding sites, SiteMap provided quantitative and graphical information in terms of site score and druggability score with properties such as hydrogen bond donor, acceptor, hydrophobic and hydrophilic regions in the predicted site. Docking studies of cytoplasmic tail region of CD80 with full length Nef revealed that CD80 may interact with the interface between the flexible N-terminal and C-terminal core domain. Potential binding sites were predicted with a good site and druggability score (>0.5; **Sup. Table 3**).

The sites of interacting regions of Nef with other cellular proteins have been previously characterized. A polyproline motif (68-78aa) present on the core domain of Nef binds to the SH3 domain of Src kinases with high (nM to μM) affinity^34^. Other than the polyproline motif within the core domain, a number of residues on the core domain are involved in multiple interactions, such as FPD_126-128_ with human thioesterase and W_61_ and L_115_ with CD4^19^. An acidic cluster (EEEE_65_) close to the core domain is required for interaction with PACS1 and controls MHC-I down-regulation^35, 36^. The unstructured regions of Nef also provide an extensive accessible surface that could be used to connect to other molecules. Since there is no prior information about the binding pattern of Nef with CD80 it was necessary to score each pose based on energy calculations. From the top ranked docked poses, the best complex with the lowest energy (−246.30 kcal/mol) was chosen as the model complex for Nef and CD80 interaction. In this predicted pose, CD80 cytoplasmic region (indicated in cyan) interacts with the Site-1 and 2 residues in the core domain of Nef (indicated in blue) (**Fig. 4c**). Based on this pose, the interacting residues were mapped (**left inset** **Fig. 4c**). The side chains of site-1 residues W_61,_ E_68_, K_99_ and R_111_ are in favourable position to interact with the N-terminus of the CD80 cytoplasmic tail. It should be noted that the residues K_99_ and R_111_ potentially make polar contact with CD80 backbone carbonyl oxygen of F_4_ and side chain hydroxyl group of Y_2_, respectively. In addition, the C-terminus of the CD80 cytoplasmic tail potentially interact with E_160_ and D_180_ residues. The side of E_160_ potentially makes salt bridge interaction with R_16_ and R_23_ of CD80 (**right inset** **Fig. 4c**,). Based on our *in-silico* predictions we chose four residues that include, 3 from Site-1 (W_61,_ K_99_ and R_111_) and one from Site-2 (E_160_) for further analysis.

### Functional validation of predicted residues mediating Nef-CD80 interaction

Single site mutant of Nef such as Nef^W61A^, Nef^K99A^, Nef^R111A^ and Nef^E160A^ were designed and purified (**Sup. Fig. 8a & b**). These mutants were tested for their affinity for CD80 peptide in the ELISA assay (**Fig. 5a**). Two mutants Nef^K99A^ and Nef^R111A^ showed a loss of affinity to the CD80 peptide, whereas, the Nef^W61A^ and Nef^E160A^ exhibited an affinity comparable to full length Nef^WT^. These mutants were also assessed for their ability to affect CD80 surface levels after delivery into APCs. Indeed, the two mutants Nef^K99A^ and Nef^R111A^ did not show any reduction in CD80 levels, while Nef^W61A^ showed slightly lesser reduction in CD80 receptors and Nef^E160A^ behaved similar to Nef^WT^ (**Fig. 5b**). Consistent with a key role for the K_99_ and R_111_ in Nef-CD80 interactions, when transduced into APCs, Nef^K99A^ and Nef^R111A^ mutants did not affect IL-2 release (**Fig. 5c**) thereby showing that T-cell activation is not compromised by these mutant Nef variants. Surprisingly, while Nef^W61A^ down modulated CD80 significantly, it did not result in a loss of T-cell activation. Since T-cell activation assay requires a minimum of 4-5 h we reasoned that the Nef^W61A^ protein delivered into the APCs may be less stable than the other isoforms for the 5 h required for this assay. Indeed, western blot analysis of the protein at 2 h versus 5 h in cell lysates shows that the level of Nef^W61A^ protein was drastically decreased after 5 h, while the levels of the other Nef protein variants remained substantial (**Sup Fig. 8c**). The other Nef^E160A^ mutant exhibited similar reduction of IL-2 release as Nef^WT^ consistent with its ability to bind CD80 peptides as well as down modulate CD80 at the APC surface (**Fig. 5b-d**).

**Fig. 5:**
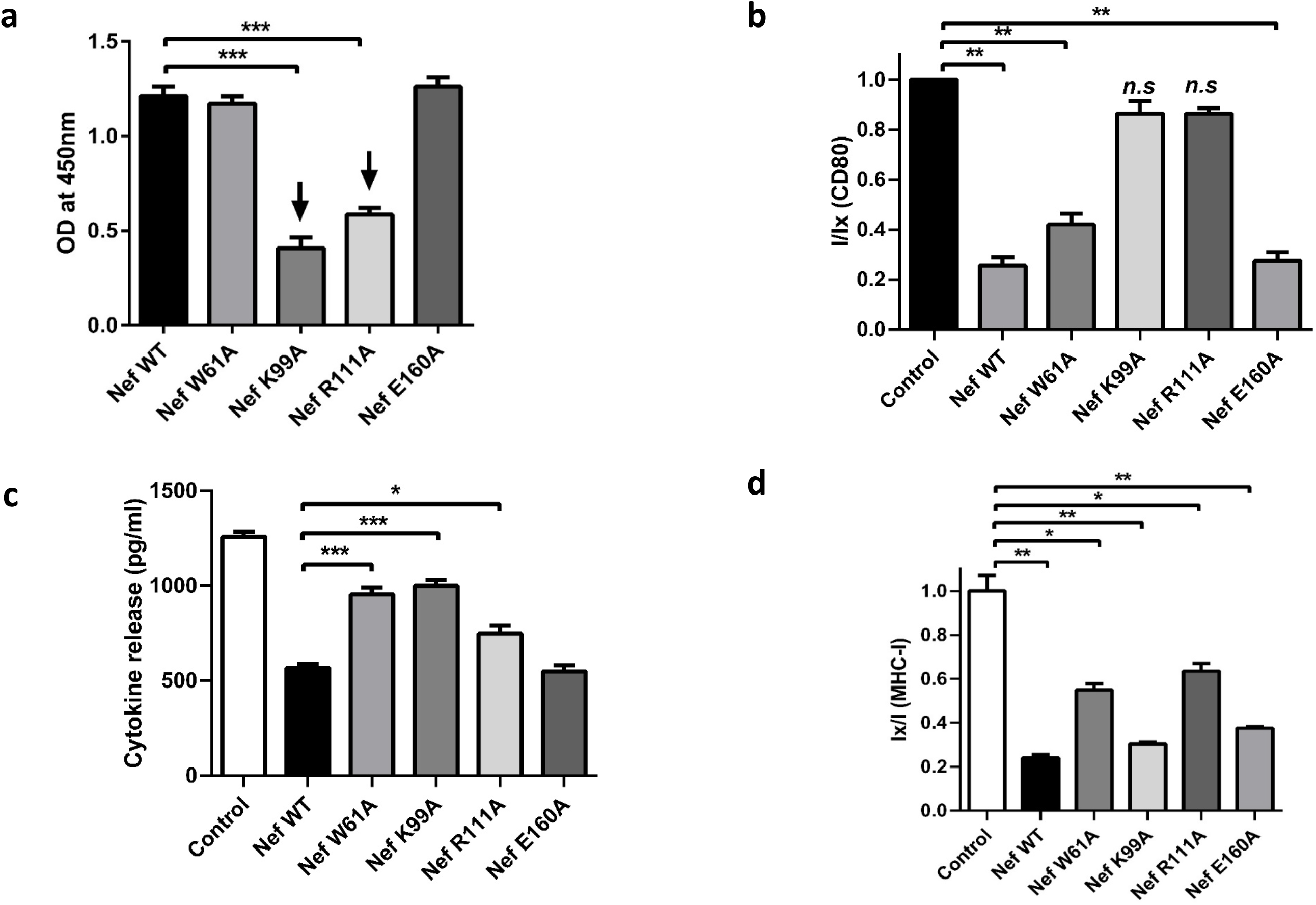
Evaluation of interaction between CD80 and Nef mutants in biochemical and cell-based assays. **(a)** Graph shows colorimetric signal of immobilized CD80 cytosolic peptide upon binding to Nef^WT^ or Nef mutants as measured by ELISA at OD450nm. Two mutants Nef^K99A^ and Nef^R111A^ showed reduced affinity to CD80 peptide **(b)** Graph shows FACS data of surface levels of CD80 receptors in RAJI cell line after delivery of Nef^WT^ or Nef mutant protein delivery. No significant down regulation seen with mutants Nef^K99A^ and Nef^R111A^ **(c)** Graph shows the levels cytokine (IL-2) released in supernatants of cells in the co-culture functional T-cell activation assay after delivery of the Nef mutants as compared to the wild type Nef protein. **(d)** Graph shows FACS data of MHC-1 levels after delivery with Nef^WT^and mutants. Nef^WT^ or mutants were delivered into RAJI cells using Chariot^TM^ delivery reagent. MHC-I was detected by flow cytometry and shown as Ix/I plots.

The two mutants Nef^W61A^ and Nef^R111A^ also did not down regulate MHC-I receptors as much as Nef^WT^; whereas Nef^K99A^ and Nef^E160A^ exhibited a similar reduction in MHC-I levels as Nef^WT^, as assessed by surface MHC-I antibody staining (**Fig. 5d**). W_61_ residue (W_57_ in Subtype B) has been previously reported to be important in CD4 down regulation and R_111_ residue (R_106_ in subtype B) is located in the oligomerization domain of Nef. Many residues of Nef have been identified that promote interaction with MHC-I, including W_57_ and R_106_ in subtype-B, NL4-3 strain, but their mutation did not hinder MHC-I down regulation^37^. However, W_61_ and R_111_ residues in subtype C, appear potentially important for Nef interaction with MHC-I, indicating subtle differences in the modulation of host proteins by different Nef variants. Nevertheless, these functional studies provide strong support to the predicted binding mode of CD80 peptide with Nef via residues K_99_ and R_111_ in Site 1 (**Fig. 4c**). Mutation in these residues leads to loss of binding capacity, resulting in the inability to down-modulate CD80 thereby restoring T-cell activation function of the transduced APCs.

### AP5 ligand binding sites in Nef full length

Considering the predictive potential of the computational model of Nef-CD80 interaction surface, the most potent binding inhibitor molecule, **AP5** was docked with Nef protein to capture its binding pattern and important residues involved in interaction. **AP5** fits nicely into the hydrophobic cavity formed by the residues W_61_, V_71,_ L_115_ and W_118_, which are part of the N-terminal loop region and the α4 helix of core domain (**Fig. 6a and inset**). The aromatic residues W_61_ and W_118_ mainly have π-π stacking interaction with aromatic ring B in **AP5**. In addition to hydrophobic interactions, the side chain of S_50_ and backbone of E_68_ form hydrogen bond interaction with amine group of the ligand. The side chain of K_99_ from α3 helix of core domain, interacts with the CF_3_ (triflouro methyl) group. Moreover, AP5 binding site overlaps with Site-1 of Nef-CD80 binding pocket, and the docking results showed that the Nef-CD80 and Nef-AP5 binding sites are overlapping with two important common residues such as W_61_ and K_99_.

**Fig. 6:**
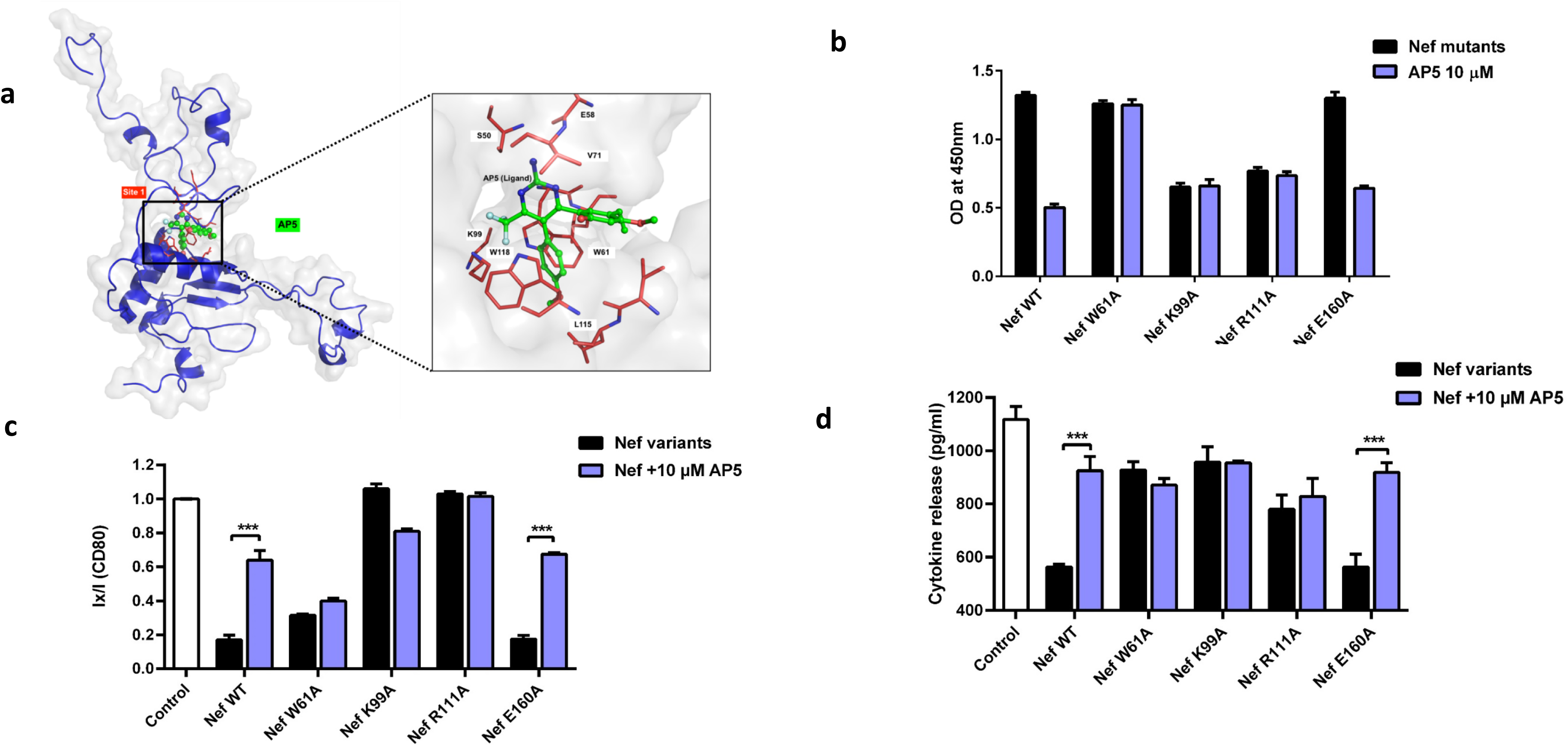
Structural and functional evaluation of the interaction between Nef and AP5. **(a)** Structural and functional evaluation of the interaction between Nef and AP5The Surface representation of HIV-1 Nef depicting **AP5** ligand (green color) binding. The binding site of **AP5** molecule overlaps with the CD80 binding site (Site-1). The inset shows the important residues for the interaction between AP5 and Nef. The non-polar residues such as W_61_, L_91_, I_109_ and L_115_ contribute to hydrophobic interactions with CF_3_. **AP5** ligand docking studies shows that the binding interactions occurs between the α4 and α5 helices along with few residues such as W_61_, E_65_ and R_111_ which are crucial for **AP5**-Nef interaction **(b)** Graph shows colorimetric signal of immobilized CD80 cytosolic peptide upon binding to Nef^WT^ or Nef mutants in the presence /absence of 10 μM AP5 as measured by ELISA at OD450 nm. (g) Graph shows surface levels of CD80 receptors in RAJI cell line after the delivery of Nef^WT^ or Nef mutant protein delivery as measured by FACS in the presence /absence of 10 μM AP5. Nef^W61A^, Nef^K99A^ and Nef^R111^ did not show any further change in CD80 levels with **AP5** addition. **(c)** Graph shows cytokine (IL-2) release in supernatants after the co-culture T-cell activation assay. RAJI cells were pre-treated with 10 μM AP5 for 1 h and then the cells were delivered with Nef mutants or wild type Nef protein for 2 h before co-culture with Jurkat T-cells for 3 h. The IL-2 levels remain unchanged with and without addition of **AP5** compound in all three mutants Nef^W61A^, Nef^K99A^ and Nef^R111^. Reduction in IL-2 seen with mutant Nef^E160A^ comparable to Nef^WT^.

Consistent with these predictions, at 10 µM **AP5** was neither able to inhibit the interaction between CD80 peptide and Nef^W61A^ mutant nor further reduce the residual interaction of Nef^K99A^ and Nef^R111A^ mutants *in vitro* (**Fig. 6b**). However, AP5 was able to displace both Nef^WT^ and Nef^E160A^ from CD80 peptides adsorbed on the ELISA plate. Thus, W_61_ is an important residue for **AP5** binding to Nef. Furthermore, in agreement with the predictions, **AP5** treatment did not result in any change in surface levels of CD80 in APCs transduced with Nef^W61A^, Nef^K99A^ and Nef^R111A^ mutant proteins (**Fig. 6c**). The levels of IL-2 release in all three mutants Nef^W61A^, Nef^K99A^ and Nef^R111A^ also remained unchanged, with the Nef^W61A^ mutant mimicking the inhibition observed with wild type Nef. **AP5** restored IL-2 release in Nef^E160A^ treated cells, comparable to Nef^WT^ (**Fig. 6d**), consistent with the inability of Nef^E160A^ to affect neither CD80 nor **AP5** binding, thereby serving as a negative control. These results predict and functionally validate the residues in the Nef protein that are important for AP5 binding.

### Hit refinement of AP series

We chose **AP5** as our starting point, given its nanomolar and micromolar potencies in biochemical and cell-based assays respectively as well as specificity to CD80. An initial hit refinement of **AP5** was conducted to understand the preliminary structure-activity relationships (SAR). New analogs were synthesized in two series by modifying the rings B and C with un/substituted aryl (heteroaryl) moieties, and without -CF_3_ group at the 6^th^ position (**Fig. 7a**). In series-1, four analogs AP(S1-S4) were prepared (**Sup. Fig. S2; Scheme-3**), where phenyl group (ring B) was placed at the 4^th^ position of ring A and varying the substitution pattern at 5^th^ position (ring B). In series-2, six analogs AP(S5-S10) were prepared by modifying both 4^th^ and 5^th^ positions (**Sup. Fig. S2**; **Scheme-4**). Apart from these, another analog **AP-S11** (**Sup. Fig. S2**; **Scheme-2**) was also synthesized where -CF_3_ was maintained at the 6^th^ position and substituted aryl rings at 4^th^ and 5^th^ positions.

**Fig. 7:**
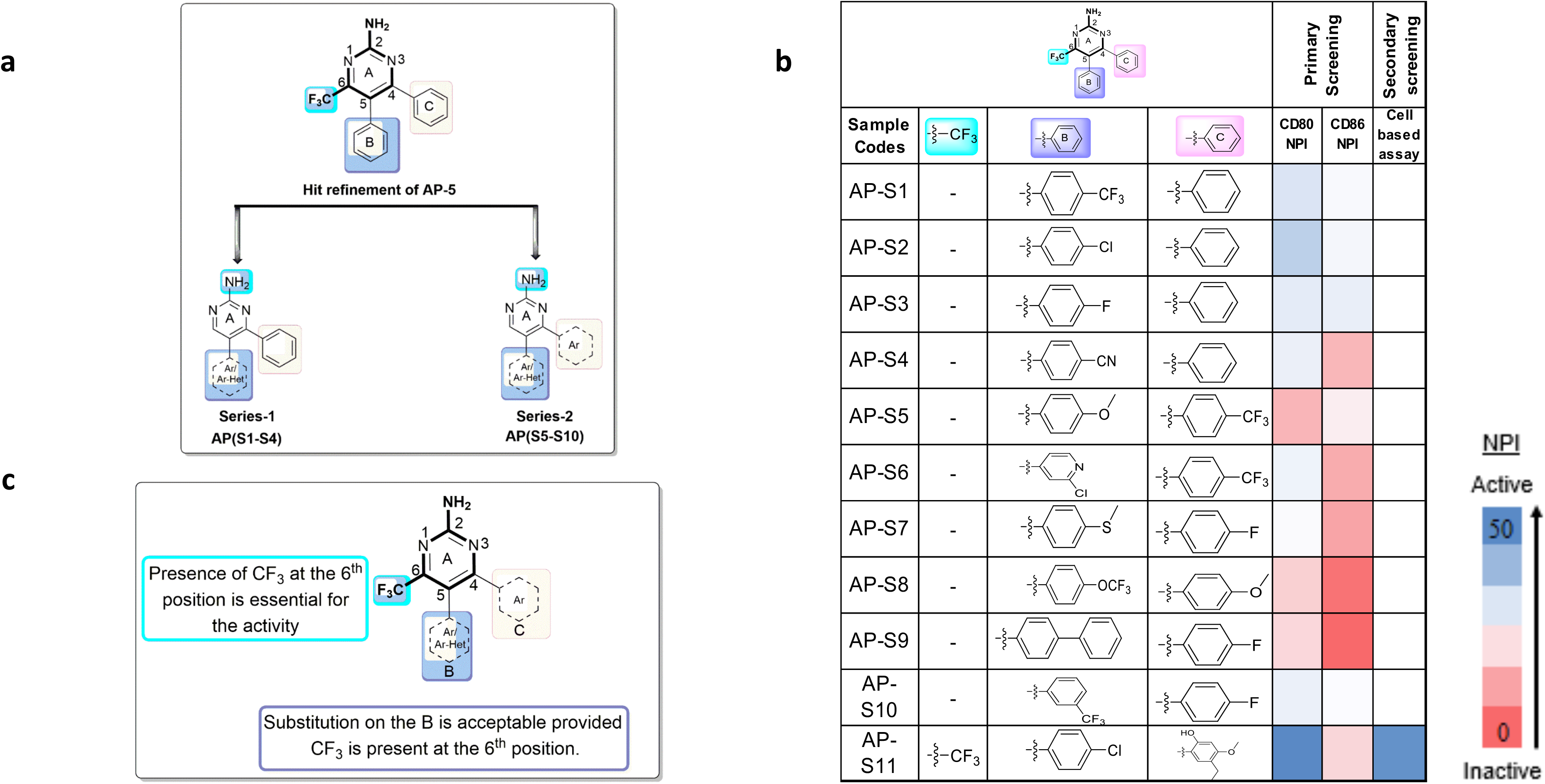
SAR and Hit refinement with AP5 as a template. **(a)** Scheme shows Medicinal chemistry approach for hit refinement of **AP5** showing two series of compounds **(b)** Summary of SAR strategy to design compounds similar to **AP5 structure** with modifications made on rings A, B and C as indicated. The synthesized molecules were evaluated for their effect on Nef-CD80/CD86 inhibition **(c)** Heat map table showing SAR with synthesized compounds. The various substitutions in rings A, B and C are indicated as well as their activity in the ELISA and cell-based assays

All the synthesized molecules were evaluated for their effect on Nef-CD80/CD86 inhibition by ELISA. The analogs **AP(S1-S10**) without -CF_3_ group at 6^th^ position of ring A didn’t show any activity (**Fig. 7b**), however, the analog **AP-S11** with CF_3_ at 6^th^ position of ring A showed the activity. These results disclosed that presence of CF_3_ group at 6^th^ position of ring A important for the activity (**Fig. 7c**). Moreover, docking studies revealed that CF_3_-group showed hydrophobic interactions with non-polar residues such as W_61_, L_91_, I_114_ and L_115_ while *in vitro* and *in vivo* experiments described above, suggested that these interactions are important for at least for the inhibition of the down modulation capacity of Nef. These studies underline the importance of the -CF_3_ group at 6^th^ position of ring A (**Fig.7a-c**). In conclusion, lead compounds with potency in nanomolar range across the cell-based assay along with acceptable solubility, permeability and pharmacokinetics parameters necessary for further drug development and chemical perturbation of the Nef-CD80 interface, have been developed.

## Discussion

HIV-Nef plays an important role in the pathogenesis of HIV infections and understanding its many functions in modifying the host cell surface has served as a focal point of HIV research^38, 39^. In earlier work we had determined that the presence of wild type Nef in virus infected cells, promotes the loss of the co-stimulatory proteins CD80/86 from the infected APC surface resulting in a loss of naïve T-cell activation^23,24,25, 40^. Here we have identified potent small molecules that disrupt the interaction of Nef with CD80 co-stimulatory receptors, and restore T-Cell activation potential of virus-infected APCs. The three lead structures identified: **AP5**, **BC5** and **PA4** belonged to three diverse scaffolds. While **PA4** and **BC5** are able to inhibit both Nef mediated CD80 as well as MHC1 down-regulation, possibly indicating different interaction points in Nef or counteracting Nef at more than one protein-protein interface. **PA4** and **BC5** molecules are leads to explore for molecular interaction promiscuity and investigating some of the multiple interactions of Nef. Since **AP5** selectively inhibits Nef-mediated CD80 down-regulation, we chose to pursue its detailed characterization in this study,

To gain an insight into the interactions of Nef with CD80, computational approaches followed by experimental validation were used to identify possible binding sites on Nef for both CD80 peptide and the small molecule inhibitor **AP5**. Due to the lack of crystals with suitable diffraction properties to provide high-resolution structures, and limited experimental information on full length Nef structure, possibly due to its inherent flexibility we adopted a computational strategy. Full length Nef was modeled by a multi-template computational approach, and their spatial conformation was validated using the constraints obtained from SAXS experiments. The interaction site for CD80 was obtained by docking of CD80 cytoplasmic tail with full length Nef model, and a number of possible binding sites for its already known protein-protein interaction sites were identified. These predictions provide key insights that could be correlated with the experimental results, identifying key residues that are involved in Nef-CD80 interaction. A caveat to be noted is that there is a limitation in finding the best biologically relevant orientation of CD80 since the docking was restricted to only the cytoplasmic tail peptide of CD80 which does not impose spatial conformation of the full length CD80 embedded in the membrane. The free cytoplasmic CD80 region fits in the energetically favorable orientation, given its steric constraints, providing verifiable insights from mutation studies. In parallel the **AP5** docking results confirms that W_61_ and K_99_ residues of Nef contributes to the interaction interface with this lead molecule. Based on the Nef-CD80 and Nef-**AP5** docking results, hotspot residues such as W_61_, K_99_ and R_111_ were mutated. This revealed that the K_99_ and R_111_ residues are crucial for CD80 binding and additionally, W_61_ plays an important role in **AP5** binding. Since the binding sites for CD80 and **AP5** are overlapping, our study provides a plausible view of the inhibitory mechanism, where AP5 interactions with Nef would prevent its ability to associate with CD80, since they compete for the same site. Ongoing efforts are aimed at improving the inhibitors with -CF_3_ group in **AP5** series and further validating these hits in *in vivo* studies.

Our results also indicate that Nef interaction with the co-stimulatory receptors CD80/86 cytoplasmic tails are distinct from reported Nef –MHC-I interactions. Nef interacts with MHC-I cytoplasmic via E_62_-_65_ and P_78_ residues^41^, although subtype C as shown here may utilize W_61_ and R_111_. While Nef-MHC-1 interactions are important, Chaudhry *et al*^24^ showed that the kinetics of Nef down-modulation of MHC molecules is slow as compared to the loss of co-stimulatory CD80/86 function. We have shown that the interaction of Nef with the co-stimulatory CD80/86 is likely to be critical for inhibiting the priming of the immune system towards naïve infections, and hence may be critical for immune evasion strategy of the virus^42, 43^. The chemical tools developed here will allow such an interrogation in suitable animal models.

Once developed into drug-like chemicals, the leads we have identified from this study would have significant impact in at least two scenarios where macrophages play an important role for HIV-1 pathogenesis. One important application is in maternal–fetal transmission and other in cases of early infection or persistent infection resulting from viral mutations. 90% of HIV infection in children is through Macrophage-tropic Maternal-fetal Transmission^44^. Nef plays a major role in vertical transmission of Macrophage-tropic HIV-1 in mother to child, where the motifs for receptor modulation were conserved in mother-to-infant *NEF* sequences. The second scenario is in early stages of infection when viral load is low, the role of Nef-mediated CD80 and CD86 down modulation from the antigen-presenting cell surface could delay the onset of T-cell responses to provide the virus with a time window sufficient for expansion. There is evidence that T-cells need co-stimulatory molecules for optimal killing of target cells, thus even in cases of established infection the removal of CD80 and CD86 from infected cell surfaces could reduce the efficiency of T-cell responses^45^. If this is the case, blockade of this function by small-molecule inhibitors of Nef-CD80/CD86-cytosolic tail interactions, could enhance effectors antiviral immunity and delay the onset of disease in both pathophysiological contexts.

Altogether, targeting this protein-protein interaction interface represents a promising new therapeutic approach to bring forth a first in class set of inhibitors to foreshorten the infection burden in HIV-1. The information gained from this integrated approach of both computational and experimental study have set the foundation for further ongoing efforts in synthesizing the next series of more drug-like inhibitors. We have demonstrated a promising chemical starting point for building chemical tools and drugs that can interfere with immunomodulatory consequences of HIV-1.

## Significance

Our study aims to develop small molecule inhibitors that disrupt the interaction interface between HIV-1 viral protein Nef and host CD80 / CD86 in Antigen Presenting Cells (APC). The disruption of this interaction will makes infected APCs more visible to the immune system, increasing cytotoxic lymphocyte activity on these HIV-infected cells, potentially leading to viral clearance from macrophage reservoirs. Here we identify and structurally characterize small molecule inhibitors that indeed disrupt the protein-protein interaction interface of Nef-CD80 and restore the T-cell activation capacity of infected APCs. These chemical tools serve as excellent starting points that may be used to interrogate the role of Nef in HIV immune evasion and contribute to first-in-class drugs for mitigating this resurgent disease

## Acknowledgments

This work was funded by the BIRAC-CRS grant (BT/CRS0045/CRS -02/12), J.C. Bose Fellowship from the Department of Science and Technology to SM, Government of India, IIIM funds CSIR Research grants GAP-2116 and HCP-0001. Center for Chemical Biology & Therapeutics fund (sanction order no. BT/PR7222/MED/31/1901/2012, dated 11.01.2013), and support from the Department of Atomic Energy (Government of India) under Project No. RTI 4006 to NCBS. We thank Drs. Satyajit Rath, Ramaswamy Subramanian and Shahid Jameel for helpful discussions. We also thank M. Hurakadli, J. Subbarao, P.Kumar for their technical assistance. The facilities provided by the Bangalore Life Science (BLiSc): Biosafety-I & II, CIFF, Instrumentation, X-ray facilities have greatly supported this work.

## Author contributions

SM, ASK, TS, PPS and RAV contributed to project initiation and funding. SM and ASK contributed to the study design of biochemical screen and cell-based assays. PPS and RAV contributed to the medicinal chemistry study design. ARV contributed to the structural biology study design and supported this work at the CCBT. AUS, APN and AM contributed to compound screening and cell-based experimental work. SS, SA, GM, KRY and PPS contributed to chemical synthesis and structure activity relationship analyses. GA and RS contributed to the molecular modeling and docking studies. SR contributed to mutant study. NK performed the SAXS studies. SM, AUS and ASK wrote the manuscript, which was reviewed by RAV and ARV. Figures were prepared by AUS, SS, PPS, GA and NK.

## STAR methods

### Materials and Reagents

96-well maxisorp ELISA plates (NUNC, cat#449824,), small molecule inhibitors (synthesized by IIIM, Jammu), DMSO (Sigma, cat#D2650), Anti-Nef Antibody (from ICGEB), DAR-HRP (Jackson Immunoresearch, cat#711-035-152), Tecan ELISA reader. The following commercially synthesized peptides (Peptron Inc., South Korea) were used: CD80 cytosolic tail peptide: PRCRERRRNERLRRESVRPV (20-mer), CD86 cytosolic tail peptide: LWKWKKKKRPRNSYKCGTNT (20-mer), CD74, a non-specific peptide: MHRRRSRSCREDQKPVMDDQRDLISNNEQL (30-mer). Nef construct was cloned at ICGEB, rF2-Nef protein (HIV-1 Subtype C) cloned into pET28 vector (Novagen, cat#69865) at NcoI and XhoI sites with 6XHis tag at C-terminal end. Nef and all Nef variant proteins were expressed in E. coli Rosetta strain containing pRARE that codes for t-RNAs corresponding to rare Arginine codons in the bacterium. The growth and/or expression medium used were Luria Broth. The antibiotics used were kanamycin (50µg/ml) for Nef plasmid selection and chloramphenicol (25µg/ul) for pRARE selection, IPTG induction, Akta FPLC –Affinity and Size exclusion chromatography for protein purification.

#### Cell lines

RAJI B lymphocytic, Burkitt’s lymphoma cell line (NIH AIDS reagent cat#ARP-9944), Jurkat T cells (ATCC, cat#TIB-152), for cell-based assays, HEK293T (ATCC CRL-3216) was used for transfection of viral clones and production of viral particles.

#### Cell culture reagents

RPMI 1640, FBS, glutamine, Penstrep (from Gibco), Chariot^TM^ delivery reagent (active motif, cat#30100) was used for protein delivery into cells. Viafect (Promega, cat#E4982) reagent was used for DNA transfection, lentiX concentrator (clontech, cat#631231), lentiblast reagent (OZ biosciences,cat#LB00500), DMEM (Gibco), WST-1 reagent (Roche), anti-CD80 and anti-CD86-biotinylated antibody (Ebioscience, cat# 13-0809-82 and 13-0869-82), SAV-APC (Ebioscience, cat#17-4317-82), anti-CD3 antibody (Biolegend, 317302), western blot materials (Biorad), Chemiluminescent reagent (Pierce, cat#32109), GE Image Quant, human IL-2 ELISA kit (Biolegend, cat#431808), Plasmid prep kit (Qiagen), biotinylated anti-human MHC-I (HLA-A, B, C) antibody (biolegend, cat#311402).

### Chemical Compound Synthesis

The 25 active hits belong to three scaffolds namely amino pyrimidine (**AP**), biaryl (heteroaryl) carbamate (**BC**) and phenoxy acetamide (**PA**) has been selected and details regarding the strategies applied for the synthesis of hits and hit optimized compounds has been provided in Scheme 1-8 (Supplementary data). The synthesis started with amino pyrimidine (AP) scaffolds, where five hits were synthesized and their synthetic strategy has been given in Scheme 1 and 2. The synthesis started with commercially available 2-chloro-5-bromopyrimidine **1** as the starting material which was treated with ethanethiol **2** afforded intermediate **3** (Scheme 1). The intermediate **3** was then subjected to Suzuki coupling with aryl boronic acid **4** provided 5-aryl pyrimidine **5**. The compound **5** was then oxidized with *m*-CPBA to corresponding sulfoxide **6** followed by nucleophilic substitution afforded key intermediate **7**, which on reaction with acetylacetone provided **AP1**. In the next attempt, the intermediate **7** on reaction with substituted aldehydes provided **AP2** and **AP3**. In another attempt, the 2-chloro-5-bromopyrimidine **1** was converted into **AP4** in two steps i) nucleophilic substitution with dimethyl amine; ii) Suzuki coupling with phenyl boronic acid. The synthesis of **AP5** required the quite different strategy and is shown in Scheme 2. The synthesis started with commercially available resorcinol **11** which was undergoes acetylation followed by reduction gave intermediate **13**. The intermediate **13** was treated with 4-methoxyphenyl acetonitrile **14** to get acylated intermediate **15** which on treatment with trifluoroacetic anhydride underwent cyclization to generate the chromone based key intermediate **16** followed by the methylation of hydroxyl group to get **17**, which on reaction with guanidine hydrochloride to afford the hit **AP5**.

For the initial hit refinement, we started synthesis with commercially available 2-aminopyrimidine **28** which on reaction with phenyl boronic acid **4b** with methodology was developed in the presence of light and K_2_S_2_O_8_ gave intermediate **29**. Intermediate **29** was brominated in the presence of *N*-bromosuccinamide (NBS) provided the intermediate **30**. The final targeted compounds **AP(S1-S4)** were synthesized in good to moderate yields from the reaction of compound **30** with substituted phenyl and heterocyclic boronic acids **4** under Suzuki conditions to provide final compounds **AP (S1-S4,** scheme-3). For series-2, sequence of synthesis began with 2-amino-4-chloropyrimidine **31** which underwent Suzuki coupling reaction with un/substituted aryl and heteroaryl boronic acids **4** to get intermediate **32** which on reaction with NBS provided intermediate **33**. The brominated intermediate **33** was subjected to Suzuki couplings with a range of aryl and heteroaryl boronic acids **4** to provide final compounds **AP (S5-S10,** Scheme-4).

In the case of biaryl (heteroaryl) carbamate (BC), seven hits were synthesized (Schemes 5-6). The synthesis started with commercially available phenylchloroformate **18** which on treatment with benzo[d]oxazole-2(3H)-thione **19** in the presence of base to give the hit **BC1** (Scheme 5). All other hit molecules from **BC3** to **BC7** and **BC10** were synthesized in the similar fashion, different substituted chloroformates **18** reacted with substituted anilines **20** to afford the targeted hits (Scheme 6).

In case of phenoxyacetamide (PA), syntheses of eight hits were accomplished as outlined in Scheme 7 and 8. Hit molecules like **PA2**, **3**, **5**, **6** and 7 (**Scheme 7**) were synthesized in two steps. i) by treating substituted anilines **20** with different chloroacetyl chloride in the presence of base at room temperature followed by; ii) coupling with substituted phenols **23**. For the synthesis of hits **PA1, 8 and 10,** the synthesized involves four steps. The substituted phenols **23** coupled with substituted 2-chloroethylacetate **24** to form intermediate **25**, which on hydrolysis gave intermediate **26**. The intermediate **26** was converted into corresponding aryl chloride **27** and then coupled with substituted anilines **20** to get the desired hits **PA1, 8** and **10** (Scheme 8).

The identified hits such as **BC2, 6 and 8**, and **PA4** and **PA9** were not synthesized and were procured in somewhat large quantities from the original commercial vendors because of the unavailability of the starting materials. These hits were characterized by using NMR and Mass spectroscopy and then were taken up for validation study.

#### Compound stock and storage

All compounds were dissolved 100% DMSO to make a 10mM stock. Multiple aliquots were prepared from mother stock to avoid multiple freeze-thaw cycles and were stored at -80°C

### Expression and Purification of recombinant Nef

HIV-1 Nef gene sequence was cloned into pET28b expression vector with antibiotic resistance to chloramphenicol (25µg/ml) and kanamycin (50µg/ml) and recombinant Nef-His tag protein was expressed in *E. coli* Rosetta strain [OD600∼0.5-0.6]. IPTG induction (0.2mM) was done for 4h at 28°C. The cells were spun down at 10000rpm for 15minand pellet stored at -80°C. The protein was purified in 20mM Tris HCl, 150mM, NaCl, 3 mM DTT, 5% glycerol, 0.2% Tween-20 in a Ni-NTA column and gel filtration chromatography on Sepharose-75pg (GE) in Akta FPLC purifier. Nef protein with single mutants W61A, K99A, R111 and E160A were designed by site-directed mutagenesis. The buffer conditions were same as full length WT-Nef.

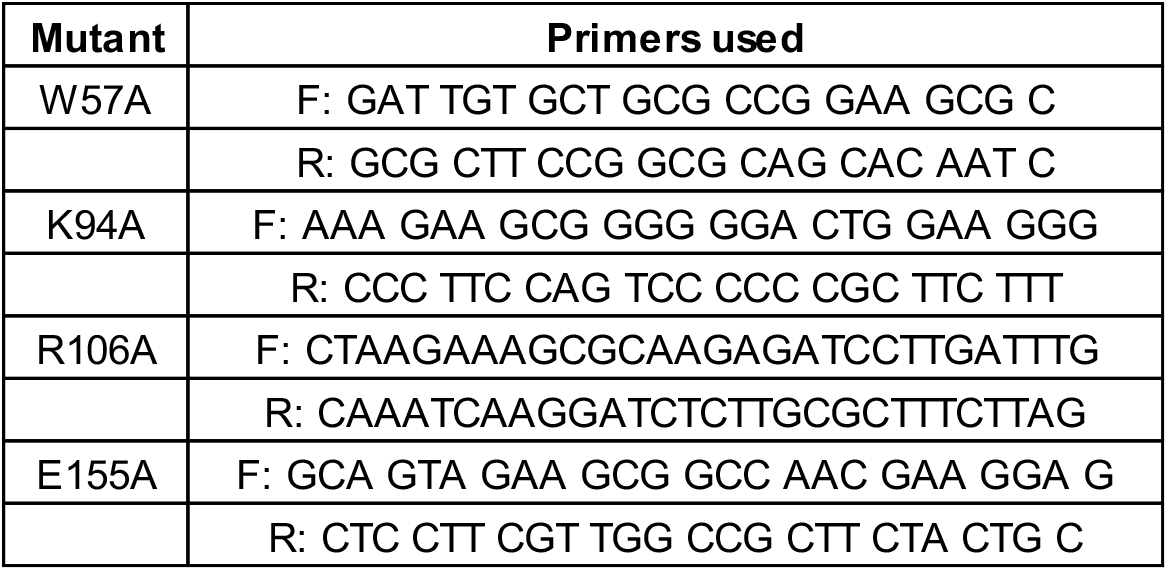

### Microscale Thermophoresis (MST)

Full length Nef was labeled with lysine NT-647-NHS fluorescent dye using the Monolith NT.115 Protein Labeling Kit (NanoTemper Technologies). A Capillary Scanning was performed to check the optimal fluorescence intensity of the labelled protein for titration with the ligand. For the direct binding assay, a final concentration of labelled 35nM Nef protein was titrated against a 16-point 2-fold serial dilution series starting from 950 µM for CD80 peptide and for CD86 a final concentration of labelled 25nM Nef protein was titrated against a 16-point 2-fold serial dilution series starting from 10 µM. The compound was titrated against the protein-peptide at16-point dilution. All samples were prepared by centrifuging at 10000 rpm for 5 min at 4°C and 10 µl of the supernatant was loaded into premium glass capillaries (NanoTemper Technologies). MST runs were performed at MST power of 60% and excitation power of 50%, using a Monolith NT.115 NanoTemper Technologies. The data was analyzed using NanoTemper analysis softwareMO. Affinity Analysis v2.2.4. Kd values were determined using T-jump and thermophoresis settings. The change in thermophoresis between each sample dilution was represented as normalized fluorescence (ΔF_norm_), which is defined as F_hot_/F_cold_, where F_cold_ is the control and F_hot_ is the experimental condition. The binding kinetics to non-fluorescent ligand causes a change in thermophoresis which is determined by area in the curve under steady-state conditions to yield a binding curve.

### Primary screening by Indirect Enzyme-linked immunosorbent assay (ELISA)

An indirect ELISA was performed to measure the interaction of Nef-CD80/CD86 and their disruption with addition of compounds.CD80, CD86 and a non-specific control peptide derived from the cytoplasmic tail of CD74 peptides were separately immobilized at 10 µM concentration onto 96-well micro-titer plate and incubated overnight at 4°C. 5% blotto was used as blocking buffer to reduce the background interference. Blotto was completely removed with PBS-Tween (0.1%) washes. 10 µM of Nef protein and 10 µM of compounds were pre-incubated at RT for 1hour and added onto peptide coated wells for 1hour incubation. Compounds were diluted in 2.5% blotto and were screened at a concentration of 10µM. After each addition step PBS-T washes were done. DMSO in 2.5% blotto as vehicle control and plate background (no coating) were used as negative control. In subsequent steps the plates were incubated with primary anti-Nef antibody and secondary antibody with HRP. The TMB substrate was added which reacts with HRP to produce a coloured product within 15minutes. The reaction was stopped with 1N sulphuric acid and the absorbance at 450nm was recorded with Tecan infinite 200 PRO plate reader. MATLAB 7.5 program was used to pick the active compounds. The program assessed the robustness of the screened plate using Z-factor (for qualifying plate) and Z score for active compounds, and Normalized percent inhibition to determine the compound activity.

### Robustness of screen

The plate controls showing a Z factor >0.5 was considered to qualify for further analysis. The activity of compounds was determined with the Normalized Percent Inhibition (NPI) and Z scores were used to assess the efficacy of the compounds in the screen: Z-score: (Ix-μD)/ σD

Normalized inhibition (NPI): ((Ix-μD)/ μD)*100

Where, IX: Intensity of triplicate x (first, second or third) represented as either A/B/C μD: Mean of negative control

σD: Standard deviation of triplicates of negative control

The compounds identified as ‘hits’ in primary screen were defined as those displaying more than 30% and 20% inhibition for CD80 and CD86 respectively and two of three repeats of a particular compound should have Z score between - 2 and -8.

#### Determination of IC_50_ for active compounds

The active compounds from primary screen were tested for dose response with 10-fold dilution for an 11-point curve from 10µM to nanomolar concentration. Compounds showing an IC_50_ in nanomolar concentrations were further tested in cell-based assay.

#### WST assay for quantification of cell viability and toxicity

50000 cells per well were seeded in a 96 well plate. 1%Triton X (control) and compounds were treated at 100 µM in triplicates in 100 µl media and incubated for the 24 hours’ timepoint. 10 µl of WST-1 reagent per well was added and incubated for 3hrs.Absorbance read at 450nm with reference 620nm. The media alone control was used to normalize all the wells. The cells alone control was the high control and triton treated cells were the low control or negative control.

The formula used to calculate the percentage toxicity:

(Average _(X)_ – Average _(high control)_ / Average _(low control)_ – Average _(high control)_)*100 Where, Average_(X)_= average OD of individual test; Average _(high control)_ = average OD of cells alone control; Average _(low control)_ = average OD of cells with 1% triton X

### Measurement of surface levels of CD80/CD86 receptor by flow cytometry

Nef protein was delivered into RAJI B cells using Chariot^TM^ protein delivery reagent according to manufacturer’s protocol (Active Motif). In brief, 1.5X10^5^ cells were layered with protein-delivery reagent complex, incubated and complete media (RPMI+10% FBS) was added. The cells were then harvested and stained with CD80 or CD86 biotinylated antibody or isotype control followed by Streptavidin-APC at 4°C and flow profile acquisition was done on Gallios Flow Cytometer (Beckman Coulter). Data was analyzed by FlowJo LLC software. Surface levels of CD80 and CD86 was calculated by normalizing raw fluorescent measurements relative to controls (Normalized Inhibition of Down-regulation). To test compounds, two concentrations 10 and 100µM were pre-treated on cells for 1 hour and then 50 μg of Nef was delivered with method described and quantified by flow cytometry.

The potential actives from the plates were selected based on the following metrics.

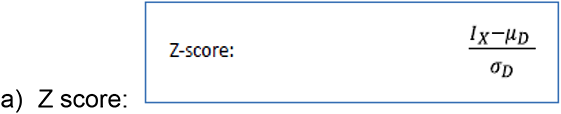

Where I_x_ is the measurement of each triplicate, μD: mean of population and σD: Standard deviation of population (excluding the positive and negative control). Z score was used to select hits reversal of CD80/86 downregulation by Nef. Compound repeats were qualified by Z score selection, with a Z score between 1.5 to -1.2. Each graph was calculated with student t-test using Graphpad prism 7.0 (*p≤0.05; **p ≤ 0.01; ***p ≤ 0.001) was used to determine the significant difference between the means of control group and treatment groups.

#### Calculation of surface CD80 and CD86 down regulation

was calculated by normalizing raw fluorescent measurements relative to controls (Normalized Inhibition of Down-regulation)

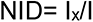

Where I_x_, is the raw measurement each triplicate and I is the mean of the measurements on the positive control.

### Virus particle generation

#### HEK293T transfection

The 4 retroviral components present in the following plasmids, pVSV-G (envelope), p-gag-pol (packaging), pMSCV-Nef-YFP (HIV-1 F2 isolate-based retroviral clones harboring nef sequence), pMSCV-YFP (control) were transfected into individual recombinant bacteria and cultured in LB for plasmid isolation (miniprep Qiagen). The plasmid concentration was checked by NanoDrop^TM^.

#### Virus particle generation

To generate recombinant retroviral particles, HEK293T were seeded into 100mm dish with 5X10^6^ cells on day 1. At 80% confluency, Viafect ^TM^ transfection reagent with 15µg of DNA vectors consisting of p-gag pol, pVSV-G and YFP or Nef-YFP in 3:1:4 ratios was layered over cells with DMEM (5% FBS). The cells are incubated for 72h. The cells were observed under fluorescence microscope for YFP expression to estimate the percentage transfection. The cell supernatant was collected and concentrated with Lenti-X^TM^ reagent (as per manufacturer’s protocol). The visible pellet was reconstituted with minimal volume of PBS (1X) to make a concentrated viral stock. The stock was titrated in 3-fold serial dilution onto HEK293T (1X10^5^) cells/ well in a 24 well plate. Polybrene (8µg/ml) was added to cells along with DMEM (5%FBS) media. After 72h, the cells were harvested and analyzed on Gallios flow cytometer. The fluorescent population was gated and analyzed against the cells alone. The infection percentage above 30% and below 0.5% is omitted. The higher titer tends to be underestimated; lower titer falls too close to the background. The average of the titer was used to calculate the viral units present in the stock using the low formula:

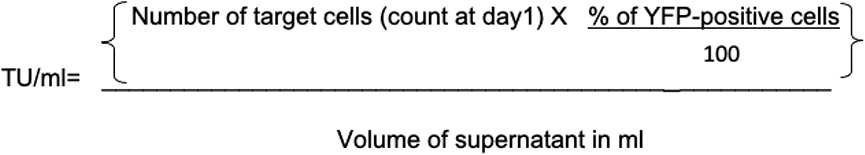

##### Functional T-cell activation Assay

RAJI cells were infected with MOI-0.02 of viral stock of control YFP virus or Nef-YFP containing virus and incubated in a 24-well cell bind plate. Additionally, LentiBlast^TM^ reagent A and B was added in 1:1 ratio to the cells and incubated for 48 to 96h. The cells showed YFP signal post infection. The cells were harvested and stained for CD80 and CD86 and analyzed by flow cytometry and the percentage down-modulation of the receptors was calculated with median values.

Virus infected RAJI-cells was co-cultured with Jurkat T cells (1:1 ratio, 2.5X10^5^ cells) in the presence of 0.06 µg/mL of anti-CD3 antibody (OKT3 clone, BioLegend®) in final volume of 200 µl and incubated for 3h for 37°C in a 96 well plate and the supernatant was harvested by centrifuging twice at 4000 rpm. IL-2 Cytokine release in the supernatant was quantified by BioLegend® kit-based ELISA method and ODs were measured in Tecan infinite 200 PRO plate reader.

#### Testing of compounds in viral assay for functional T-cell activation assay

Pre-treatment with compounds **AP5, PA4** and **BC5** at 1, 10 and 100 µM concentrations was performed for 24h before Nef viral transduction in RAJI cells. Vehicle control cells were pre-treated with 0.5% DMSO. Post-treatment of compounds for 24h was done after 96h of viral infection.

### Staining of surface MHC-I antibody

RAJI cells were pre-treated with compounds **AP5, PA4** and **BC5** at 100 µM and 50 µg Nef was delivered with Chariot^TM^ reagent. The cells were stained for receptors with anti-human MHC-I biotinylated (HLA-A, B, C, clone W6/32) BioLegend® antibody and secondary streptavidin APC and was run on Gallios flow cytometer and the surface levels MHC-I was quantified. The percentage down regulation recovery of MHC-I receptors was analyzed.

### Modelling of full-length Nef

To build a full-length Nef protein, multi-template modelling approach was performed, where more than one experimentally determined structure was utilized for building the model. For the N-terminal part, NMR structure of Nef anchor domain(1QA5) and for core domain, the NMR structure of HIV-1 Nef (2NEF: A) and X-ray structure HIV-1 NEF protein, in complex with engineered HCK SH3 domain (3RBB: A) are used as templates. Among the three templates, major structural information is acquired from 2NEF structure which covers maximum region of core domain. The tool MODELLER7 (version 9v8) was used to obtain the full-length Nef model. After modeling, the lowest energy state structure was further energy minimized through SYBYL (Version 7.1) (Tripos Associates Inc.) and validated using PROCHECK^46^

### Deciphering the residues necessary for Nef - CD80 interaction by docking studies

The modeled full-length Nef protein is utilized for examining the potential-ligand association site using the SiteMap^c^ tool in Schrödinger software. SiteMap identifies potential peptide/ligand binding sites considering van der Waals forces and hydrogen donor ⁄ acceptor characteristics. SiteScore is the most important property generated by SiteMap, proven to be effective at identifying possible binding sites in 3D structure. The prediction of the binding site is based on set of properties such as size of the site, degrees of enclosure by the protein and exposure to solvent, tightness with which the site points interact with the receptor, hydrophobic and hydrophilic character of the site. The sitemap predictions are useful in identifying the possible binding sites of CD80/CD86 cytoplasmic tails.

Further, docking studies were performed to identify the interactions of Nef protein with co-stimulatory molecules CD80/CD86. This was achieved by modelling of cytoplasmic regions of CD80/CD86 using I-TASSER server and protein-peptide docking using BioLuminate module in Schrödinger software. Though numerous information about the Nef interaction sites with other cell surface receptors are available, the peptide is docked by blind docking approach where no guidance about residues, that could potentially participate in interaction with CD80 was provided to the program. The best docked pose is selected by energy minimization followed by implicit solvent based energy calculations.

The protein-peptide docking resulted in 30 best poses of Receptor-ligand complexes. The predicted poses are ranked based on the maximum number of occurrences of that particular pose. Since there is no prior information about the binding pattern of Nef with CD80/CD86, it is necessary to score each pose based on energy calculations. The association of the protein-peptide complex is estimated by an automated mechanism of Multi-Ligand Bimolecular Association with Energetics (eMBrAcE) (MacroModel, version 9.6, Schrödinger, LLC, New York, NY, 2008). The best identified Nef-CD80 binding pose analysed further for important amino acids involved in the non-bonded interactions which is contributing for binding. In order to validate the predicted binding mode, best binding small molecule **AP5** was docked with the full-length Nef model. This was basically achieved by selection of a centroid point from the predicted Nef-CD80 binding site using Glide docking protocol from Schrodinger software. The top poses of the docked complexes were further examined for the non-bonded interactions and best docked score.

### SAXS data collection and analysis

SAXS-data of the apo HIV1-Nef was measured with the BIOSAXS-1000 small-angle X-ray scattering with Kratky camera system, installed on a Rigaku microfocus X-ray generator (1.5418 Å wavelength). The purified HIV1-Nef at 1, 3 and 5 mg/ml concentrations were used with buffer containing 50 mM Tris/HCl, pH 7.5, 200 mM NaCl in a sample volume of 60 µl inside a vacuum tight quartz capillary subjected X-rays at 25 °C. The data was collected for 30 min and for each measurement a total of six frames at 5 min intervals were recorded. Corresponding to each protein sample, data were collected for a buffer under identical experimental conditions, providing a background scattering curve. The data was then tested for possible radiation damage by comparing the six data frames and no changes were observed. The scattering of the buffer was subtracted from the scattering of the sample. All the data processing steps were performed using the program package PRIMUS^47^. The experimental data obtained for all protein samples were analyzed for aggregation using the Guinier region. The forward scattering I(0) and the radius of the gyration, Rg were computed using the Guinier approximation assuming that at very small angles (q<1.3/Rg) the intensity is represented as I(q)= I(0)exp(-(qRg)2/3). These parameters were also computed from the extended scattering patterns using the indirect transform package GNOM, which provides the distance distribution function P(r) of the maximum particle dimension, Dmax as well as the radius of gyration, Rg, qualitative particle motion was inferred by plotting the scattering patterns in the normalized Kratky plot ((qRg)2(I(q)/I(0)) vs qRg). Ab initio low-resolution models of the proteins were built by the program DAMMIF^48^ considering low angle data (q<2nm^-1^). Ten independent ab initio reconstructions were performed for each protein and then averaged using DAMAVER^49^. Superimposition between ab initio reconstruction and atomic model was performed using the software SUPCOMB^50^.

### Statistical Analysis

Z factor was used for qualifying the plate and t-test was used for determining the significant data. The plots were prepared using Graph Pad Prism Ver6.0.

## Supplementary Information

### Supplementary figure legends

**Fig. S1: a)** Sequence of Subtype B and C with overlapping residues and important residues highlighted (in red) **(b).** Sequence of full length CD80 and CD86, where the 20-mer cytoplasmic peptide used for screening assays are highlighted (in cyan) **(c)** Schematic of ELISA procedure in microwell plate where the CD80 and CD86 peptides were immobilized. The Nef protein was incubated with the peptide and their interaction was detected by anti-Nef antibody and secondary antibody with HRP. The colorimetric signal was quantified with TMB reducing the HRP substrate

**Fig. S2:** Detailed chemical synthesis of the hits belongs to the AP, PA and BC scaffolds. **Note: Details of Characterization of Synthesized compounds and intermediates will be included in the full submission**.

**Fig. S3:** Regression graph showing correlation with replicate data R^2^ values for hit compounds in biochemical screen, where pIC_50_ is the negative log of IC_50_ expressed in molar units.

**Fig. S4:** Graph shows cytotoxicity profile with treatment of compounds. RAJI cell line was treated at the highest concentration of 100 μM for 24 h and the supernatant collected was estimated for WST assay.

**Fig. S5:** FACS histogram shows surface staining of CD80 and CD86 with APC-tagged specific CD80 or CD86 antibody. Colour representation: Unstained Cells (in red), Isotype controls -IgG1k for CD80; IgG2b for CD86 (in blue dotted line), cells with Vehicle control Chariot reagent showing CD80 or CD86 surface expression (in orange), Protein delivered (in green). The reduction in CD80/CD86 levels are seen with Nef protein after its delivery into RAJI cells in 2h incubation period. The surface levels of CD80 did not change with the delivery of 100 µg of Ovalbumin and β-lactoglobulin proteins with Chariot^TM^ reagent.

**Fig. S6: Schematic of the reterovirus used for infection assays.** Distribution of regulatory elements in the HIV-1. (A) Schematic depiction of the HIV-1 genome containing accessory *vpr*, *vpu*, and *nef* genes. (B) Schematic depiction of reteroviral vector with Nef transgene. where Env protein used in this reterovirus is VsVg from pMLV, and Vpr, Vif and Vpu are not present in this vector.

**Fig. S7: a)** SAXS-patterns of full length HIV1-Nef at three different concentrations 1, 3 and 5 mg/ml are shown. The Guinier plots at low angles appeared linear and showed no aggregation (shown in inset) **(b)** The Rg, Dmax and molecular mass of the full length HIV1-Nef suggest the progressive increase in the Rg, Dmax and molecular mass values with increasing concentration of Nef. The deviation from a typical bell-shaped profile depicts an inherent structural flexibility of Nef Protein **(c)** The averaged ab initio model (surface representation) overlay with the crystal structure of folded C-terminal core (cartoon representation).

**Fig. S8: a)** SDS-PAGE run with purified Nef proteins. Nef wildtype and mutants show bands at ∼29kDa **(b)** FPLC purification profile of Nef proteins purified (c) Western blot showing Nef WT or mutant Nef protein delivered in RAJI cells post 5 h. Band intensity of Nef^W61A^ protein reduced after 5 h when compared to 2 h incubation, indicating degradation of protein.

### Supplementary tables

**Suppl. Table 1:** a) Table shows 20 compounds selected from screen showing IC_50_s in sub-micromolar ranges. Of these top 10 were chosen based on ease of synthesis, efficacy and their cytotoxicity.

**Suppl. Table 2:** Table showing the SAXS derived parameters such as Guinier Rg, Realspace Rg and Dmax for the full-length Nef in solution

**Suppl. Table. 3**: Table showing the residue information about the CD80 peptide docked predicted sites on Nef from SiteMap program. The important residues involved in protein-protein interactions are highlighted.

## Abbreviations

Nef: (Negative Regulatory Factor)
MHC: (Major Histocompatibility Complex)
CD80/86: (Cluster of Differentiation 80 and 86)
APC: (Antigen Presenting Cells)
AP: (Aminopyrimidine)
PA: (Phenoxyacetamide)
BC: (Biaryl (heteroaryl) carbamate)
ART: (antiretroviral therapy)
NNRTI: (non-nucleoside inhibitors)
HAART: (Highly Active Anti-Retroviral Therapy)
BnAb: (Broadly neutralizing antibodies antibody)
LTNPs: (Long-Term Non-Progressors)
MST: (Micro-Scale Thermophoresis)
WST-1: (Water Soluble Tetrazolium-1)
SAXS: (Small Angle X-ray scattering)

## Graphical Abstract

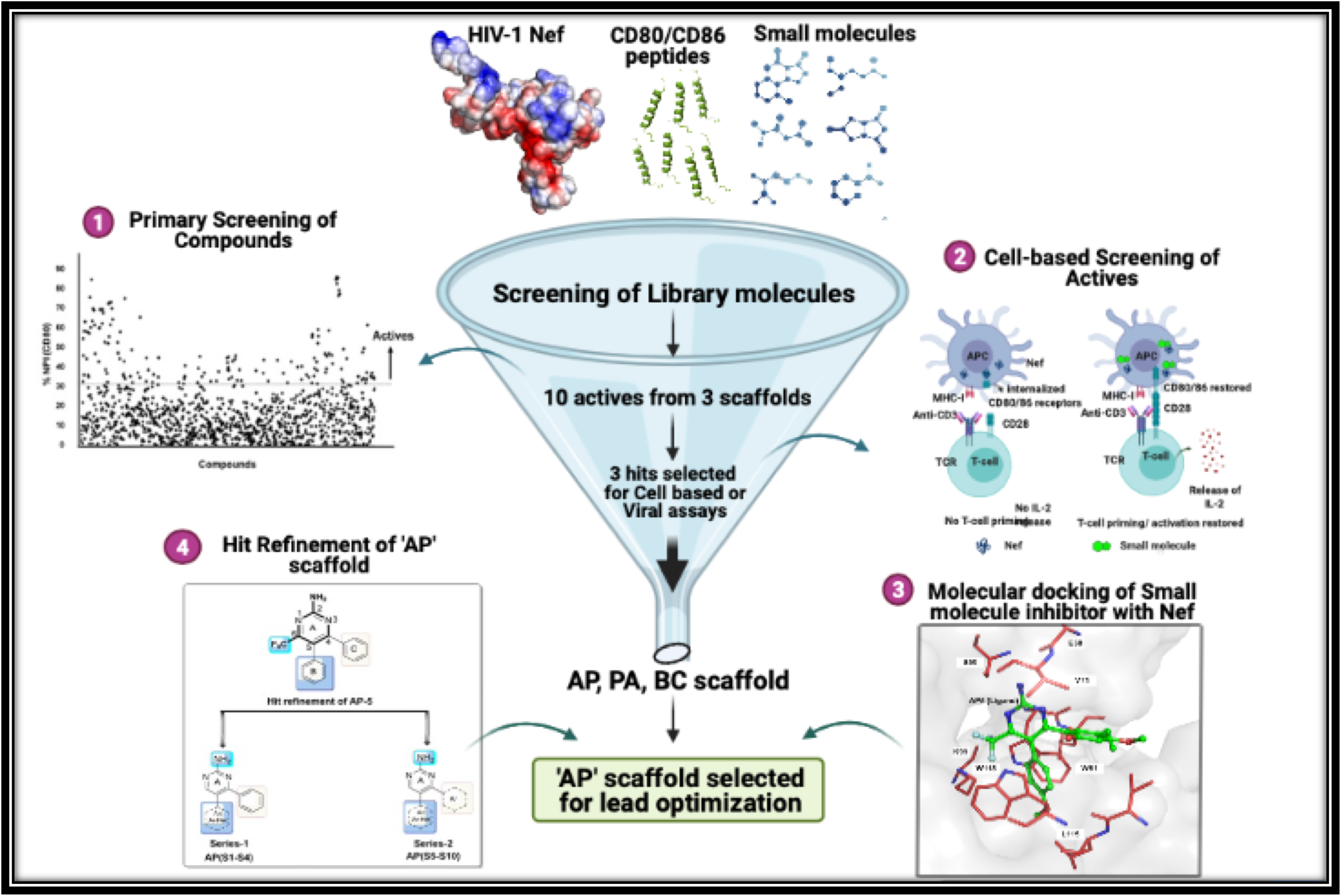

